# Aldo-Keto Reductase Family 1 Member A1 (AKR1A1) Deficiency Exacerbates Alcohol-Induced Hepatic Oxidative Stress, Inflammation, Steatosis, and Fibrosis

**DOI:** 10.1101/2022.12.07.519420

**Authors:** Ying-Wei Lan, Wan-Ru Chen, Chih-Ching Yen, Kowit-Yu Chong, Ying-Cheng Chen, Hueng-Chuen Fan, Ming-Shan Chen, Chuan-Mu Chen

## Abstract

**Background:** Alcohol-associated liver disease (ALD) covers a wide range of hepatic lesions that depend on the amount and duration of alcohol consumption, from early and reversible conditions to hepatic steatosis and severe lesions, including steatohepatitis and alcoholic fibrosis, to irreversible cirrhosis. AKR1A1, an aldo-keto reductase family member, participates in the detoxification of alcohol-derived acetaldehyde, but its role in ALD remains unclear. In this study, we studied the role of AKR1A1 in the development of ALD using *Akr1a1*^-/-^ knockout mice and palmitic acid/oleic acid (P/O) plus ethanol-treated AML12 hepatocyte cells.

**Methods:** Levels of AKR1A1 were measured in mice fed with the Lieber-DeCarli diet containing 5% alcohol (alcohol-fed, AF) or control liquid diet (pair-fed, PF). The effects of AKR1A1 on the liver function, inflammation, oxidative stress, lipid accumulation, and fibrosis were assessed in AF-induced *Akr1a1*^-/-^ and ICR control mice.

**Results:** Data showed that *AF-Akr1a1^-/-^* mice exhibited an exacerbation of liver injury and increased gene and protein levels of inflammatory mediators, oxidative stress, lipid accumulation, and fibrosis, whilst decreased expression of antioxidant enzymes in their livers than the AF-ICR mice. Therefore, loss of AKR1A1 can activate 4-HNE/p53 signaling to modulate ROS and antioxidant balance, increase lipid peroxidation, fatty acid synthesis and lipid droplet formation, reduced fatty acid β-oxidation, and elevated proinflammatory and fibrotic mediator, eventually exacerbate the ALD. In *in vitro* study, we further demonstrated that knockdown of *Akrlal* aggravated the effects of alcohol plus P/O-induced oxidative stress and steatosis, LPS-stimulated inflammation, and TGF-β1-induced fibrosis in AML12 hepatocyte cells.

**Conclusion:** our results revealed that AKR1A1 exerts protective effects on alcohol-induced liver injury, steatosis, and fibrosis, possibly by regulating the 4-HNE-p53 signaling pathway.

## Introduction

Alcohol-associated liver disease (ALD), one of the most common chronic liver diseases, is caused approximately 2 million deaths per year worldwide [1]. ALD is caused by excessive alcohol consumption and led to the development of hepatitis, steatosis, cirrhosis, and some liver cancers. The 5-year mortality rate of patients with alcohol-associated hepatitis who have related cirrhosis is more than 65% [2]. The imbalance between pro-oxidant generation and antioxidant defenses, increased gut-derived endotoxin, activation of immune response, liver cell apoptosis and necrosis, and fibrosis are currently proposed as the major elements in the progression of ALD [3–5]. Therefore, in-depth mechanism research on the pathogenesis of alcohol-induced liver diseases is needed to better prevent or treat the occurrence of ALD.

The liver contains multiple enzymes involved in alcohol metabolism. Two key enzymes, alcohol dehydrogenase (ADH) and cytochrome P450 2E1 (CYP2E1) are abundant alcohol-oxidizing enzymes in the liver and are mainly responsible for converting alcohol into acetaldehyde, which can directly cause damage brought upon by alcohol [6,7]. A third enzyme, aldo-keto reductase family 1 member A1 (AKR1A1), can detoxify toxic aldehydes in an NADPH-dependent manner. These toxic aldehydes have been identified from a variety of metabolically bioactive compounds in diabetes and cancers. The AKR1A1 enzyme is also involved in ascorbic acid synthesis [8,9]. AKR1A1 levels are abundant in the liver and can prevent carbon tetrachloride-[10], acetaminophen-[11–13] and N-nitrosodiethylamine-[14] induced hepatic injury.

Because AKR1A1 mediates the denitrosylation of S-nitrosylated glutathione and coenzyme A in mammals to regulate nitric oxide-based metabolic signaling, the anti oxidative capacity of AKR1A1 knockout mice has been shown to be significantly increased, which may protect against thioacetamide-induced hepatic injury [15] and ischemia/reperfusion (I/R)-induced acute kidney injury [16]. However, the function of AKR1A1 in the alcoholic liver disease remains unclear. In this study, we investigated the role of AKR1A1 in the development of experimental chronic alcoholic liver disease by administering an alcoholic diet in *Akr1a1*^-/-^ knockout mouse model and also explored the mechanism of AKR1A1 signaling pathway in palmitic acid/oleic acid (P/O) plus ethanol-treated AML12 hepatic cells.

## Materials and methods

### Animals and chronic alcohol feeding model

Male ICR mice (WT) aged 8 weeks were purchased from Lasco (Taipei, Taiwan).

Mice genetically altered for AKR1A1 knockout (*Akr1a1*^-/-^ ICR background) had previously been produced in our laboratory [17]. The chronic alcohol feeding procedure followed the National Institute on Alcohol Abuse and Alcoholism (NIAAA) model as previously described [18]. Eight-week-old male ICR and *Akr1a1*^-/-^ mice fed with Lieber-DeCarli liquid diets containing alcohol (alcohol-fed; AF) or isocaloric maltodextrin control liquid diet (pair-fed; PF) for 8 weeks. Glass liquid diet feeding tubes containing both liquid diets were used and the concentration of alcohol in the liquid diet gradually increased to 1%, 2%, and 4% (vol/vol) ethanol every 2 days and finally to 5% ethanol in the liquid diet for 7 weeks. Both liquid diets were changed every day, and the consumed volumes were recorded. At the end of 8 weeks of feeding, cardiac puncture was performed to obtain blood samples from isoflurane-anesthetized mice, and then the epididymal fat and livers were collected and stored at −80°C.

### Biochemical analysis

Mouse blood was collected into a microtainer serum separator gel tube (BD Microtainer #365967; BD Biosciences, Ann Arbor, MI, USA) and allowed to clot for 30 min at room temperature before centrifugation for 2 min at 12000 x g. The levels of aspartate aminotransferase (AST) and alanine aminotransferase (ALT) were measured using an IDEXX VetTest Chemistry Analyzer (IDEXX Laboratories Inc., Westbrook, ME, USA). Hepatic tissues (~100 mg) were lysed with appropriate homogenizing buffer to detect the hepatic triglyceride contents (BioVision, Milpitas, CA, USA), GSSG/GSH (Cayman, Ann Arbor, MI, USA), and malondialdehyde (MDA; Cayman) by commercial ELISA kits according to the manufacturer’s protocol.

### RNA isolation and quantitative real-time reverse transcription–PCR (qRT–PCR)

Total RNA was isolated from hepatic tissue or cell lines using the Presto^™^ DNA/RNA/Protein Extraction Kit (Geneaid Biotech Ltd., Taipei, Taiwan). RNA was reverse transcribed into cDNA using an MMLV Reverse Transcription kit (Protech, Sparks, NV, USA) as previously described [19]. qRT**–**PCR was performed using qPCRBIO SyGreen Mix (Protech) and the QuantStudio^™^ 6 Pro Real-Time PCR System (Applied Biosystems/Thermo Fisher Scientific Inc., Waltham, MA, USA). Relative gene expression was determined by the ΔΔCq method, where Cq is the threshold cycle. The relative mRNA expression levels were normalized to the mRNA level of the reference β-actin gene. Sequences of the gene-specific primers used are listed in Table S1.

### Western blot analysis

Precipitation of protein in the flow-through from the RNA Binding Step with ice-cold acetone according to the manufacturer’s protocol. Western blot analysis was performed as described previously [20]. Briefly, 50 μg of protein was separated via SDS-polyacrylamide gel electrophoresis (SDS**–**PAGE; Bio**–**Rad Laboratories, Inc., Hercules, CA, USA) and transferred to polyvinylidene difluoride (PVDF) membranes (Merck Millipore, Burlington, MA, USA). Then, the membranes were blocked with 5% skimmed milk powder (Anchor Inc., Caribbean Park, VIC, Australia) in Tris-buffered saline with 0.1% Tween 20 (TBST) for 1□h at room temperature and probed overnight with the primary antibodies at 4°C, as listed in Table S2. After the Western Lightning ECL Plus substrate (Merck Millipore) was applied, the chemiluminescent signal was detected by the use of a LAS-4000 chemiluminescence detection device (Fujifilm, Valhalla, NY, USA). The optical density of each protein was quantified by ImageJ software and normalized to that of the β-actin band.

### Assessments of liver histology

For histological analysis, one lobe of liver tissue was fixed with 10% formalin overnight and paraffin embedded. Liver tissue was sectioned and stained according to a standard laboratory protocol with hematoxylin-eosin (H&E), Masson’s trichrome (MT), and Sirius red as previously described [21]. Steatosis was evaluated by oil red O (ORO) staining of 10 μm sections from snap-frozen OCT-embedded liver tissue. Then, the cells were stained with ORO according to standard protocols as previously described [22]. Formalin-fixed, paraffin-embedded liver sections were deparaffinized, rehydrated, and incubated with a 1:200 dilution of anti-4-HNE and anti-AKR1A1 primary antibody overnight at 4°C primary antibody along with the Novolink^™^ Max Polymer Detection System (Leica Biosystems, Wetzlar, Hesse, Germany) and DAB substrate (3,30-diaminobenzidine, Leica Biosystems). Hematoxylin used for nuclear staining before mounting. Representative images were captured by using an Olympus IX71 microscope with an AxioCam MRc camera. The positive areas of collagen, Sirius red, and ORO were quantified by using ImageJ software with the IHC toolbox plug-in (National Institutes of Health, Bethesda, MD, USA).

### Cell culture and treatment

The murine hepatocyte, AML12 cell line (BCRC-60326), was purchased from Bioresource Collection and Research Center (Hsinchu, Taiwan). AML12 cells were maintained in Dulbecco’s modified Eagle’s medium/Ham’s Nutrient Mixture F-12 (DMEM/F12; Thermo Fisher Inc., Waltham, MA, USA) supplemented with 10% fetal bovine serum (Thermo Fisher Inc.), 1X Insulin-Transferrin-Selenium (ITS-G; Thermo Fisher Inc.), 40 ng/ml dexamethasone (Sigma-Aldrich, St. Louis, MO, USA) and 1% penicillin/streptomycin (Thermo Fisher Inc.) [20].

To establish a cellular model of alcoholic-induced liver steatosis, cells were treated with P/O consisted of a 2:1 mixture of palmitic acid and oleic acid with 1.8% fatty acid-free BSA and at a final concentration of 1 mM for 72 h. To mimic the alcoholic-induced liver steatosis, ethanol (200 mM) was simultaneously exposed P/O-treated AML12. After treatment, the cells were collected for the oil red O (ORO) staining and qRT–PCR analysis. To establish a cellular model of liver fibrosis, cells were treated with TGF-β1 (5 ng/ml) for 48 h, then cells were collected for the qRT–PCR analysis. To mimic liver inflammation, AML12 cells treated with LPS (1 μg/ml) for 24 h was established, then cells were collected for the qRT–PCR analysis.

### Lentiviral transduction for knockdown

Lentiviral particles for control shRNA (shCtrl) or shRNA target to *Akr1a1* gene (TRCN0000042319; 5’-GCCGATGGAACTGTCAGATAT-3’) were produced and purchased from National RNAi Core Facility (Taipei, Taiwan). AML12 cells were transduced at a multiplicity of infection (MOI) of 10 in the presence of polybrene (8 μg/ml) and incubated at 37°C for 24 h, followed by selection with puromycin (8 μg/ml) for subsequent experiments.

### Oil red O staining

AML12 cells were seeded at a density of 1×10^5^ cells/well in 12 well plates and were cultured overnight to confluency. Cells were treated with P/O plus ethanol for 72 h. Then cells were wash with DPBS twice, fixed with 10% formalin at room temperature for 30 min, rinsed with 60% isopropanol for 5 min, and stained with ORO at room temperature for 20 min to detect lipid droplets in cells. After pictures were captured by AxioCam MRc camera, the cells were washed with 60% isopropanol twice and the dye was extracted by 100% isopropanol. The intracellular lipid content was measured at wavelength of 492 nm.

### Statistical analysis

Data are presented as the means ± SD and were analyzed using GraphPad Prism 8 (GraphPad Software, San Diego, CA, USA). The statistical significance of differences between two groups was analyzed using Student’s t-test (two-tailed). One-way analysis of variance analysis (ANOVA), followed by Dunnett’s post-hoc test, was used for multiple-group comparisons. For all analyses, data were considered significant at a *p* value < 0.05.

## Results

### Chronic alcohol feeding increases AKR1A1 expression in ICR mouse liver, without liver steatosis

The C57BL/6 mice exhibited high susceptibility to developing alcohol-induced fatty liver disease (Fig. S1A) and intracellular lipid accumulation (Fig. S1B) than ICR (WT) mice. The eight-week liquid alcohol-fed (AF)-C57BL/6 mice exhibited a decreased survival rate (Fig. S1C) and body weight (Fig. S1D) and an increased liver weight and liver-to-body-weight ratio (Fig. S1D) compared with the PF-C57BL/6 mice. However, no significant differences in body and liver weight, survival rate, or lipid accumulation in the liver could be found in the ICR (WT) mice between the PF and AF groups (Fig. S1).

To explore the possible mechanism of no significant change in the ICR (WT) mice during chronic alcohol feeding (Fig. 1A and 1B), we focused on liver AKR1A1 signaling, and the expression levels of AKR1A1 in the liver were examined by immunohistochemistry and Western blotting. The level of AKR1A1 was clearly increased in the livers of the AF-WT mice compared to the PF-WT mice (Fig. 1C), and Western blotting showed the same expression pattern (Fig. 1D). These results suggest that chronic alcohol feeding increased liver AKR1A1 levels in the ICR (WT) mice.

**Figure 1.**
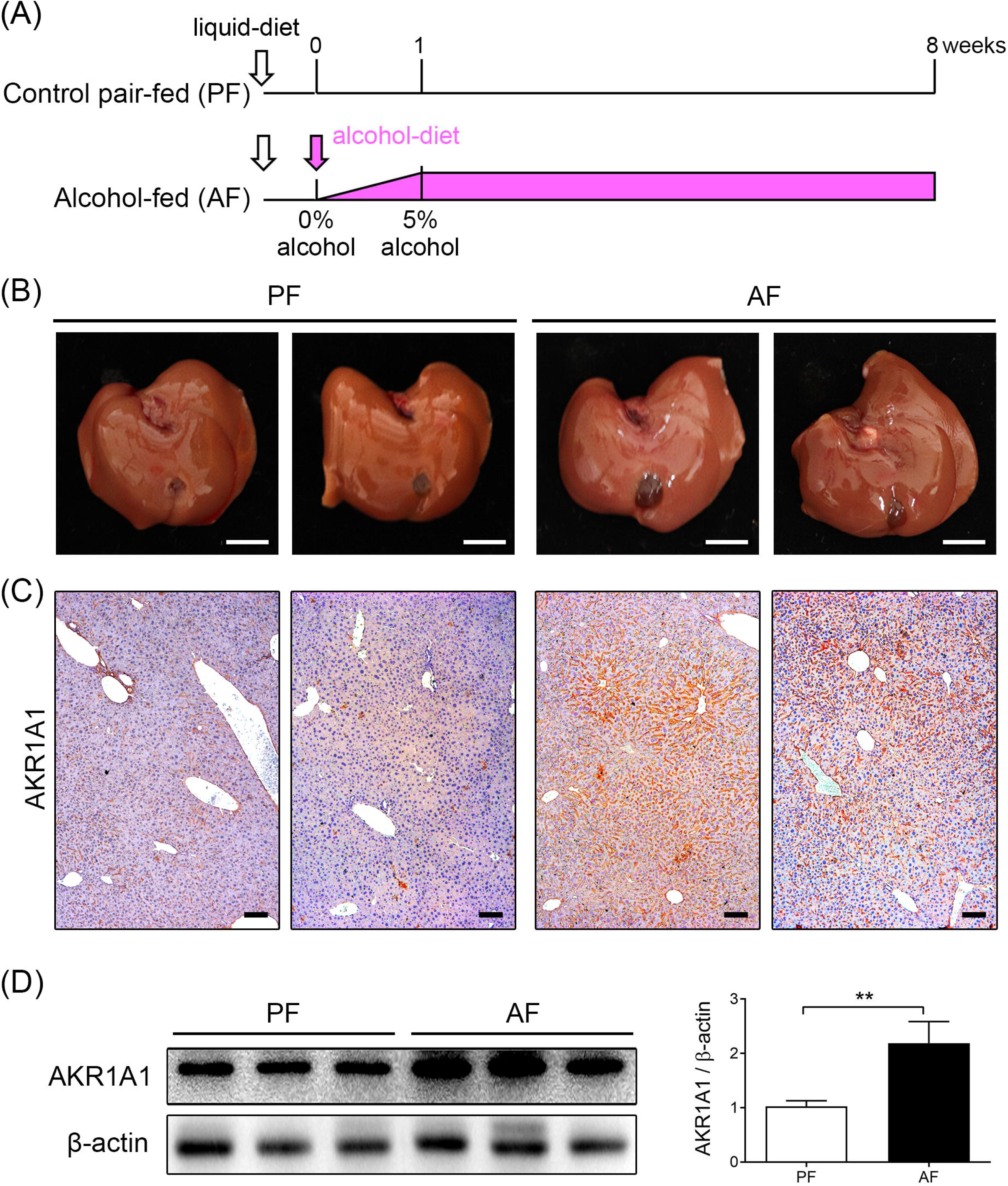
Alcohol exposure increases AKR1A1 expression in the livers of ICR mice fed a Lieber-DeCarli liquid diet containing 5% alcohol for 8 weeks. (**A**) Schedule of alcohol and control diet regimens. (**B**) Gross liver appearance (scale bars = 1 cm) and (**C**) liver sections of PF- or AF-mice after 8 weeks were subjected to IHC staining for the AKR1A1 protein (scale bars= 100 μm). (**D**) Western blotting was performed to detect the expression of AKR1A1 in the liver (left panel), and a histogram shows the densitometric quantification of AKR1A1 protein normalized to β-actin levels and expressed relative to the PF group (right panel). Data are expressed as means ± SD (n=5). \*\**p* < 0.01. PF, pair-fed; AF, alcohol-fed; IHC, immunohistochemistry.

### *Akr1a1* knockout exacerbates alcohol-induced liver injury and inflammation

To investigate the possible role of AKR1A1 in ALD, *Akr1a1*^-/-^ mice were exposed to AF. As shown in Fig. 2A, a 40% death rate was observed in *Akr1a1*^-/-^ mice after 4 weeks of alcohol feeding, and a 60% death rate was reached after 8 weeks of alcohol feeding, whereas no mouse deaths were observed in the PF-*Akr1a1*^-/-^, PF-WT, and AF-WT groups. Successful knockout of *Akr1a1* was examined by Western blotting and IHC staining, and no AKR1A1 expression was detected in the livers of either the AF- or PF-*Akr1a1*^-/-^ mice (Fig. 2, B and F). Eight weeks of alcohol feeding in the *Akr1a1*^-/-^ mice not only significantly reduced body weight, epididymal fat weight and fat-to-body-weight ratio but also increased liver weight and liver-to-body-weight ratio compared to the *PF-Akr1a1^-/-^* mice (Table S3). Furthermore, the AF-*Akr1a1*^-/-^ mice showed 10-fold increases in serum ALT levels (Fig. 2C) and 20-fold increases in AST levels (Fig. 2D), indicating liver dysfunction; a 3-fold increase in hepatic triglyceride levels promoted hepatic steatosis compared to the *PF-Akr1a1^-/-^* mice (Fig. 2E). In the AF-*Akr1a1*^-/-^ mice, H&E staining of liver sections revealed an increased number of fat droplets and few inflammatory cell aggregates (Fig. 2G). Western blotting (Fig. 2H) and qRT–PCR (Fig. 2I) results showed that the protein and mRNA expression levels of the possible proinflammatory cytokines TNF-α and IL-1β were significantly increased in the livers of the PF-*Akr1a1*^-/-^ mice compared to the PF-WT mice. Furthermore, the AF-*Akr1a1*^-/-^ mice showed higher expression levels of inflammatory cytokines than the PF-*Akr1a1*^-/-^ mice.

**Figure 2.**
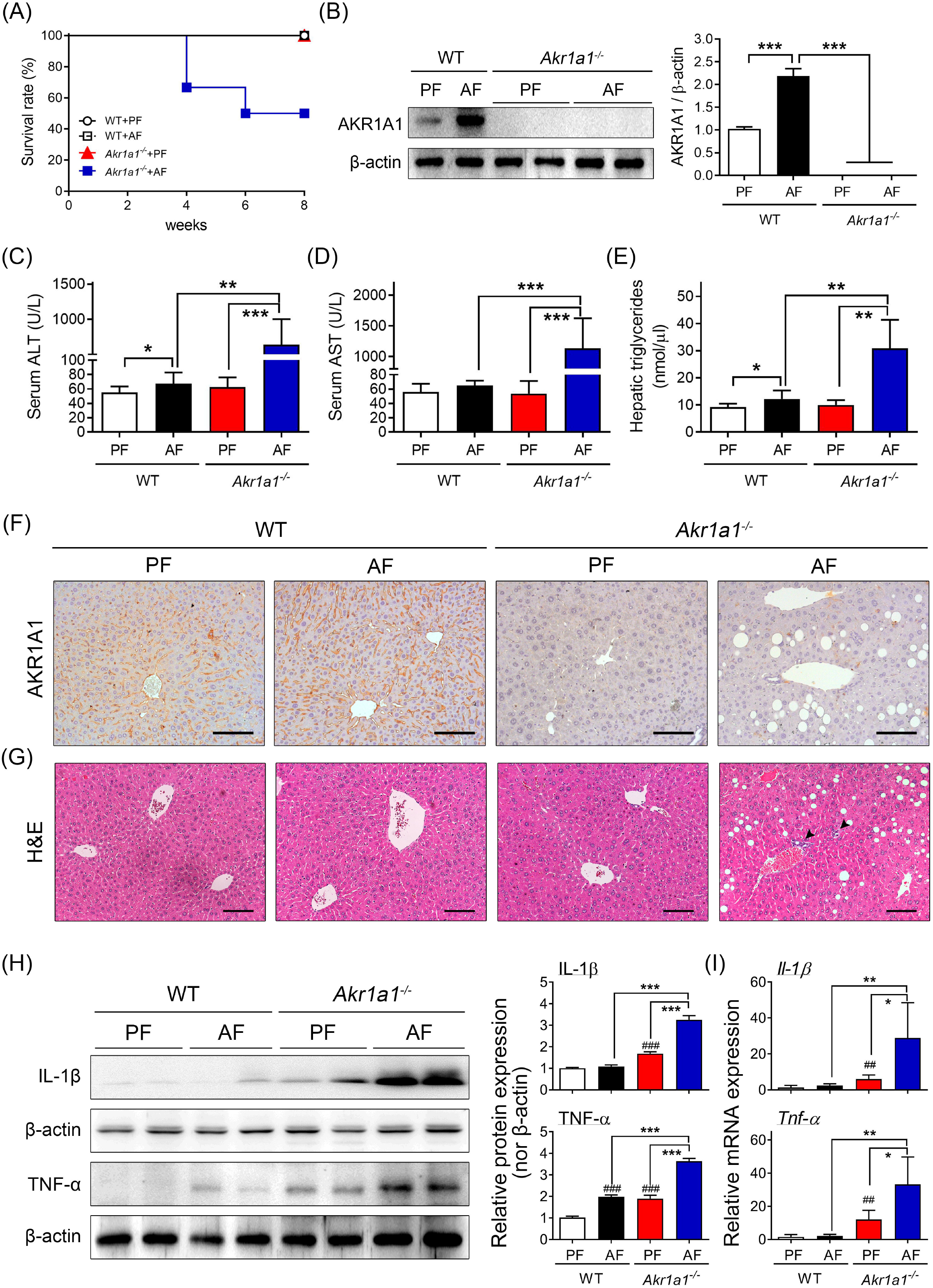
*Akr1a1* knockout exacerbates alcohol-induced liver injury and inflammation. WT and *Akr1a1*^-/-^ mice were fed a Lieber-DeCarli liquid diet containing alcohol (AF) or maltodextrin (PF) for 8 weeks. (**A**) Comparison of survival rates between four groups, WT+PF, WT+AF, *Akr1a1*^-/-^+PF, and *Akr1a1*^-/-^+AF. (**B**) Western blot analysis of AKR1A1 expression in the liver (left panel) and a histogram showing the quantitative densitometry of AKR1A1 protein normalized to β-actin levels and expressed relative to the WT PF group (right panel). (**C**) Serum ALT and (**D**) AST levels and (**E**) hepatic triglyceride contents were detected. (**F**) Liver sections were stained with AKR1A1 antibody, and (**G**) H&E staining showed the hepatic architecture with the central vein (CV). The arrowhead indicates mononuclear cell infiltration surrounding the CV. Scale bar□=□ 100□μm. (**H**) Western blot analysis of IL-1β and TNF-α protein levels in the liver (left panel) and a histogram in the right panel showing the quantitative densitometry protein normalized to β-actin levels and expressed relative to the WT PF group. (**I**) mRNA expression levels of *I1-1β* and *Tnf-α* were determined by qRT–PCR. Values were normalized to the β-actin gene and expressed in relation to the WT PF group. Data are represented as the means□±□SD; n ≥5 per group. \**p* < 0.05, \*\**p* < 0.01, and \*\*\**p* < 0.001; ^##^*p* < 0.01, and ^###^*p* < 0.001 compared with the PF-WT group. WT, wild type; PF, pair-fed; AF, alcohol-fed; ALT, alanine aminotransferase; AST, aspartate aminotransferase; H&E, hematoxylin and eosin; CV, central vein.

### *Akr1a1* knockout exacerbates alcohol-induced liver oxidative stress

To assess the effects of AKR1A1 on alcohol-induced pro-oxidant and antioxidant status and cell apoptosis-related factors, the mRNA expression levels of a pro-oxidant, *Nox-2* (Fig. 3A), and those of apoptosis-related factors, *Casp3* and *Casp8* (Fig. 3, D and E), were markedly upregulated in the liver tissue of the AF-*Akr1a1*^-/-^ mice compared to the PF-*Akr1a1*^-/-^ mice. In contrast, the antioxidant mRNAs *Sod-1* and *Nqo-1* were markedly decreased (Fig. 3, B and C) in the livers of the AF-*Akr1a1*^-/-^ mice compared to the PF-*Akr1a1*^-/-^ mice. The activities of the liver antioxidant enzymes GSH were significantly decreased in the AF-*Akr1a1*^-/-^ mice compared with the PF-*Akr1a1*^-/-^ mice (Fig. 3G). The oxidized form of glutathione, GSSG, showed no significant difference between each group (Fig. 3H), but the GSSG/GSH ratio signifying oxidative stress was also significantly increased in the AF-*Akr1a1*^-/-^ mice compared with the PF-*Akr1a1*^-/-^ mice (Fig. 3I).

**Figure 3.**
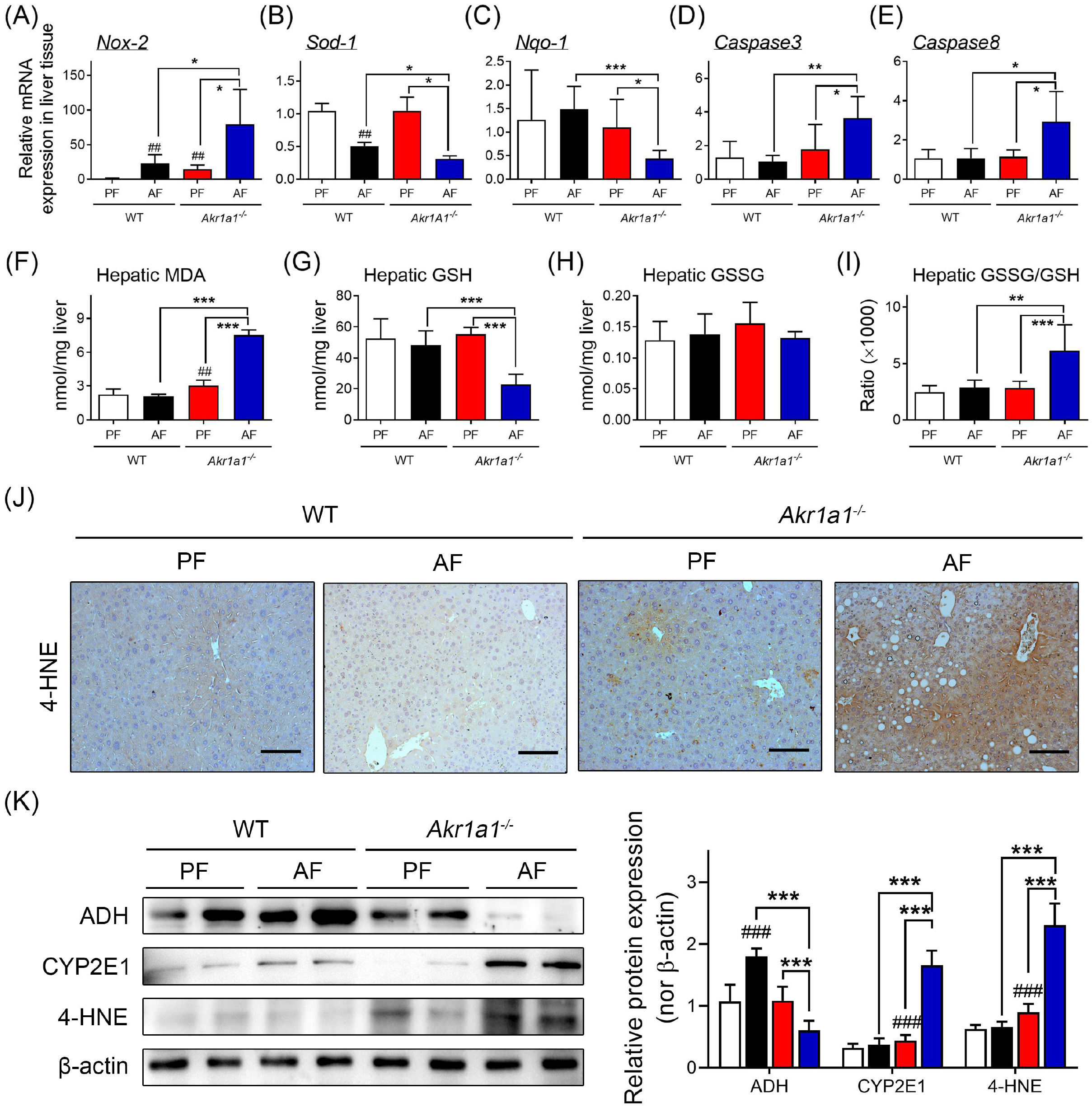
*Akr1a1* knockout exacerbates alcohol-induced hepatocyte oxidative damage. Liver tissues were collected from WT and *Akr1a1*^-/-^ mice fed AF or PF for 8 weeks. mRNA expression levels of the oxidative burst gene *Nox-2* (**A**), the antioxidant factors *Sod-1* (**B**) and *Nqo-1*(**C**), and the apoptosis-related genes *Casp3* (**D**) and *Casp8* (**E**) were determined by qRT–PCR. Values were normalized to the β-actin gene and expressed in relation to the WT PF group. (**F**) MDA levels, (**G**) GSH levels, (**H**) GSSG levels, and (**I**) GSSG/GSH ratio in the liver were determined by the indicated methods. (**J**) IHC staining of the 4-HNE protein to detect lipid peroxidation levels in liver sections. (**K**) Western blot analysis of ADH, CYP2E1, and 4-HNE expression in the liver (left panel) and a histogram on the right panel showing the quantitative densitometry of protein normalized to β-actin levels. Data are represented as the means □±□ SD; n ≥□5 per group. \**p* < 0.05, \*\**p* < 0.01, and \*\*\**p* < 0.001; ##*p* < 0. 01, and ^###^*p* < 0. 001 compared with the PF-WT group. WT, wild type; PF, pair-fed; AF, alcohol-fed.

Hepatic MDA and 4-hydroxynonenal (4-HNE) are reliable markers of lipid peroxidation, representing the oxidative stress status. The results showed that the concentration of hepatic MDA was elevated in the livers of the PF-*Akr1a1*^-/-^ mice compared to the PF-WT mice (*p* < 0.01) but was markedly increased in the liver tissue of the AF-*Akr1a1*^-/-^ mice compared to the PF-*Akr1a1*^-/-^ mice (*p* < 0.001; Fig. 3F). Similar to the MDA expression pattern, the liver 4-HNE level was examined by IHC staining and Western blotting (Fig. 3, J and K). Elevation of liver 4-HNE was detected in the PF-*Akr1a1*^-/-^ mice compared to the PF-WT mice but was markedly increased in the liver tissue of the AF-*Akr1a1*^-/-^ mice compared to the PF-*Akr1a1*^-/-^ mice (*p* < 0.001). On the other hand, liver ADH levels significantly increased to convert alcohol to acetaldehyde in the AF-WT mice, but very low ADH expression was detected in the livers of the AF-*Akr1a1*^-/-^ mice (Fig. 3K).

### *Akr1a1* knockout exacerbates alcohol-induced liver steatosis

The accumulation of droplet fat in the liver can be increased by fatty acid uptake, fatty acid synthesis increment, and fatty acid oxidation decrement. To investigate the role of AKR1A1 in alcohol-induced hepatic steatosis, the mRNA levels of genes related to lipid metabolism were measured. After eight weeks of alcohol feeding, the mRNA levels of the fatty acid uptake-related genes *Cd36, Vldlr, Fatp1*, and *Lpl* (Fig. 4A) and fatty acid synthesis-related genes *Acaca, Fasn, Srebp1*, and *Lipin1* (Fig. 4C) were markedly upregulated, but the fatty acid oxidation-related genes *Cpt1α, Acox1*, and *Ppara* (Fig. 4B) were markedly downregulated in the liver tissue of the AF-*Akr1a1*^-/-^ mice compared to the PF-*Akr1a1*^-/-^ and AF-WT mice. Liver lipid accumulation was further confirmed by ORO staining (Fig. 4E). We found that knockout of *Akr1a1* promoted alcohol-induced liver lipid droplet accumulation compared to the PF-*Akr1a1^-^* mice (Fig. 4, D and E). Western blot results also showed that the expression levels of adipose differentiation-related protein (ADRP), SREBP1, FASN, and phospho-ACC were significantly increased in the AF-*Akr1a1*^-/-^ mice compared to the PF-*Akr1a1*^-/-^ mice (Fig. 4F). These data suggested that *Akr1a1* knockout exacerbates alcohol-induced steatosis in the livers of mice.

**Figure 4.**
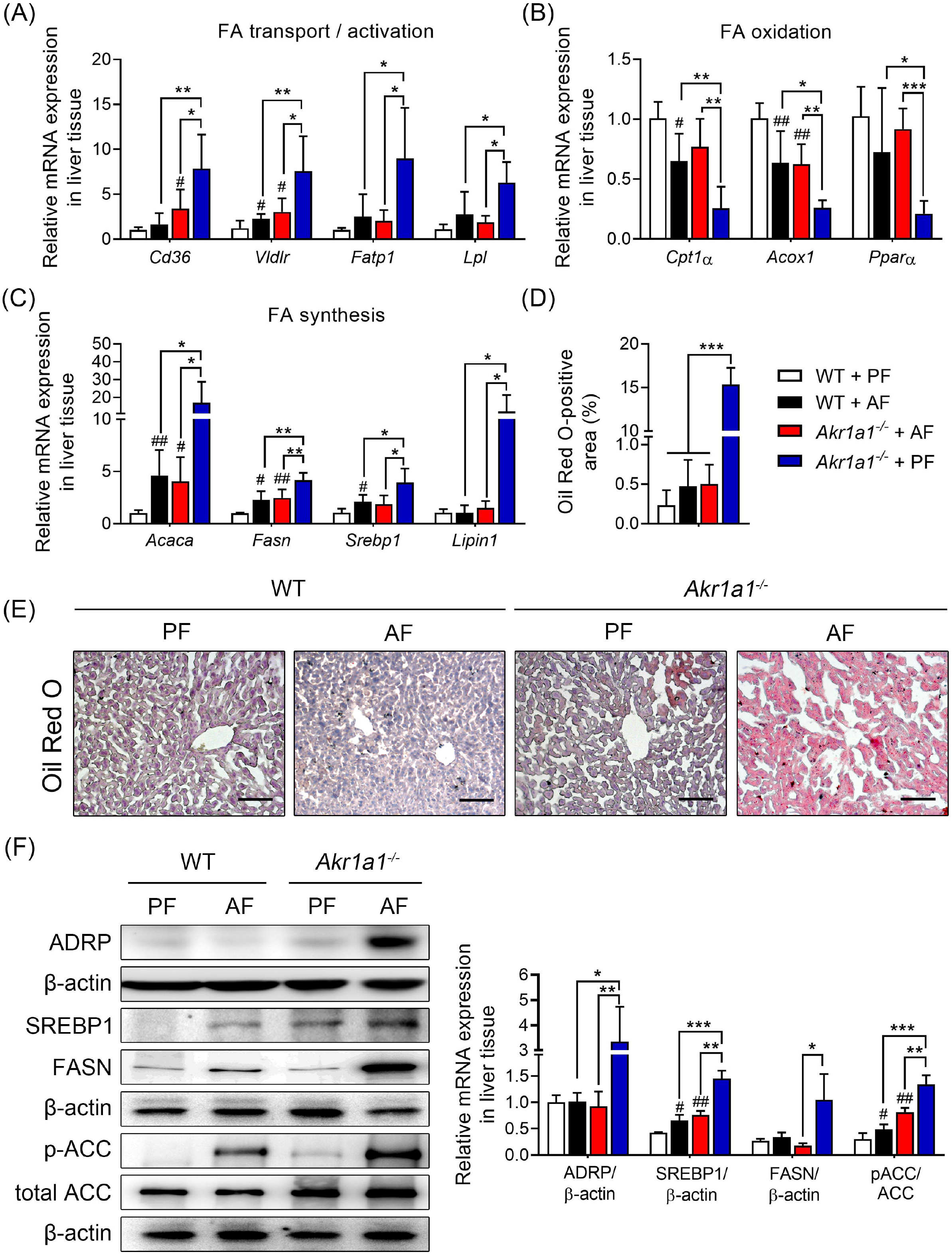
*Akr1a1* knockout exacerbates alcohol-induced lipid metabolism and liver steatosis. Liver tissues were collected from WT and *Akr1a1*^-/-^ mice fed AF or PF for 8 weeks. qRT–PCR was performed to detect the mRNA expression levels of (**A**) fatty acid transport/activation, (**B**) fatty acid oxidation, and (**C**) fatty acid synthesis. Values were normalized to the β-actin gene and expressed in relation to the WT PF group. (**E**) ORO staining of the liver lipids in frozen liver sections, and (**D**) the percentage of ORO-stained area was measured. (**F**) Western blot analysis of ADRP, SREBP1, FASN, and pACC/ACC expression in the liver (left panel) and a histogram on the right panel showing the quantitative densitometry of protein normalized to β-actin levels. Data are represented as the means±SD; n ≥5 per group. \**p* < 0.05, \*\**p* < 0.01, and \*\*\**p* < 0.001; #*p* < 0.05, and ##*p* < 0. 01 compared with the PF-WT group. WT, wild type; PF, pair-fed; AF, alcohol-fed; ORO, oil red O.

### *Akr1a1* knockout exacerbates alcohol-induced liver fibrosis

Long-term and stepwise feeding with an alcohol-containing liquid diet may promote fatty liver, leading toward liver fibrosis. Masson’s trichrome and Sirius red staining of liver tissue sections revealed a higher collagen fibrous area in the AF-*Akr1a1*^-/-^ mice than in the PF-*Akr1a1*^-/-^ mice (*p* < 0.0001; Fig. 5, A and B). Measurements by qRT–PCR showed that the mRNA levels of *Tgf-β1, Collagen I, Ctgf*, and *α-Sma* were increased in the livers of the PF-*Akr1a1*^-/-^ mice compared to the PF-WT mice (*p* < 0.05); moreover, the AF-*Akr1a1*^-/-^ mice showed markedly higher expression levels of these fibrotic factor genes than the PF-*Akr1a1*^-/-^ mice (*p* < 0.01; Fig. 5, C–F). In view of the upregulation of oxidative, steatosis, inflammatory and fibrotic mediators, possible mediators and liver p53 levels were investigated by Western blot analysis. Phosphorylated p53 showed increased levels in the *AF-Akr1a1^-/-^* mice (*p* < 0.05; Fig. 5G).

**Figure 5.**
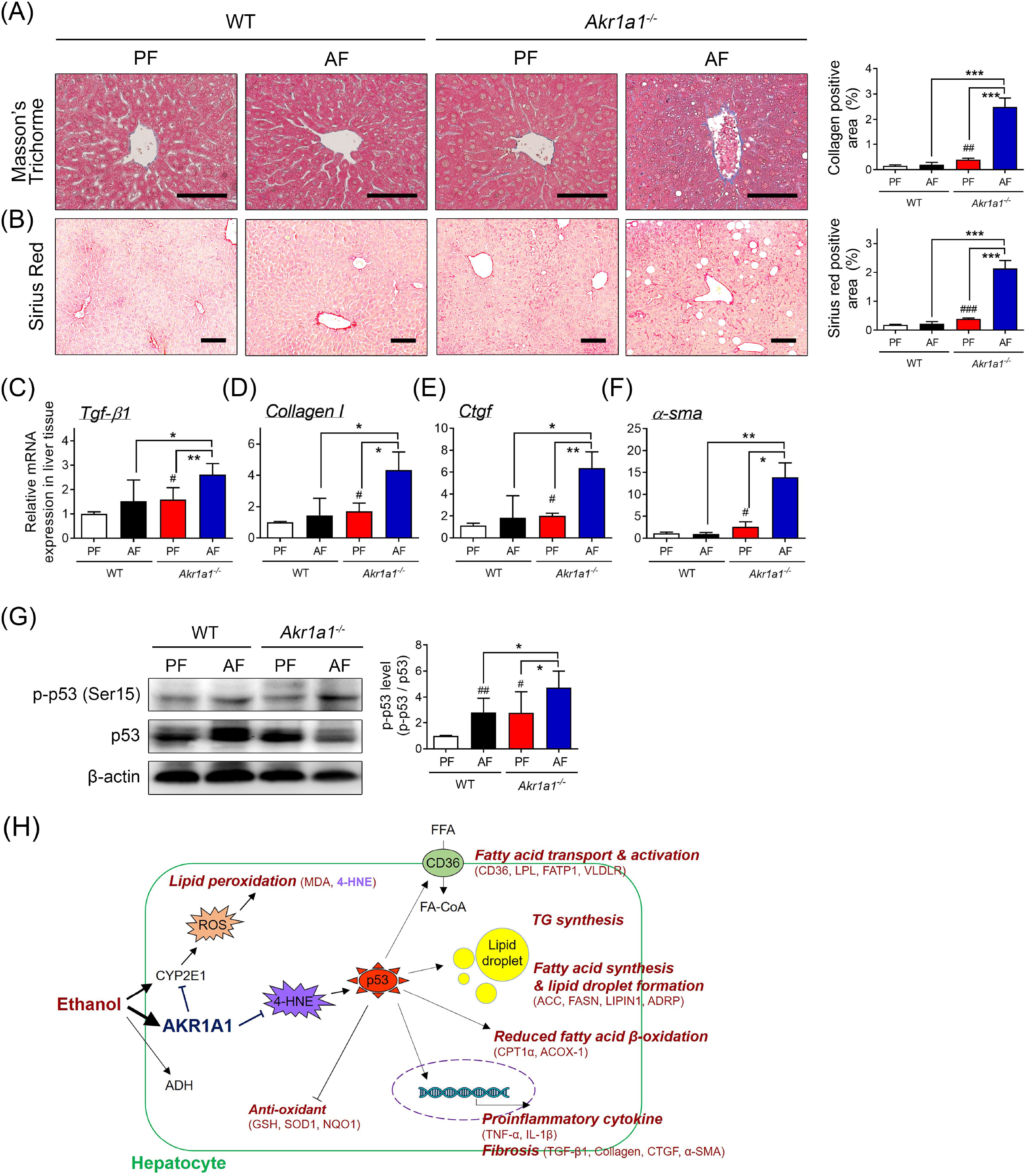
*Akr1a1* knockout exacerbates alcohol-induced liver fibrosis. Liver tissues were collected from WT and *Akr1a1*^-/-^ mice fed AF or PF for 8 weeks. (**A**) Masson’s trichrome and (**B**) Sirius red-stained liver sections from each group and the percentage of positively stained area was measured (right panel). Scale bar□=□100□μm. qRT-PCR was performed to analyze the mRNA expression levels of fibrotic-related genes, (**C**) *Tgf-β1*, (**D**) *Colla1*, (**E**) *Ctgf*, and (**F**) *α-sma*. Values were normalized to the β-actin gene and expressed in relation to the WT PF group. (**G**) Western blot analysis of p-p53 (Ser15) and p53 expression in the liver (left panel) and a histogram in the right panel showing the quantitative densitometry of protein p-p53 normalized to its p53 levels and expressed relative to the WT PF group. (**H**) Possible mechanism by which AKR1A1 regulates the development of ALD in the liver. Alcohol exposure elevated AKR1A1 levels to inhibit the expression of CYP2E1 and 4-HNE and inactivated the p53 signaling pathway to regulate ALD. Data are represented as the means □±□SD; n ≥□5 per group. \**p* < 0.05, and \*\**p* < 0.01; #*p* < 0.05 compared with the PF-WT group.

These data suggested that *Akr1a1* knockout exacerbates ALD, possibly through inhibition of the 4-HNE/p53 signaling pathway in the livers of mice (Figure 5H).

### Knockdown of *Akr1a1* in hepatocyte cells aggravates oxidative stress, steatosis, inflammation, and fibrosis

To clarify the role of AKR1A1 in pro-oxidant/antioxidant regulation, lipid droplet accumulation, inflammatory cytokine expression and ECM production, a stable cell line of *Akr1a1*-knockdown with short hairpin RNAs (shRNAs) in the AML12 cell linewas used (Fig. 6). The expression of AKR1A1 was decreased by approximately 80% at both the protein (Fig. 6A) and mRNA (Fig. 6B) levels in AML12 cells treated with AKR1A1 shRNA lentiviral particles. ORO staining cell images and quantitative data on the amount of extracted ORO staining (Fig. 6, C and D) showed that alcohol plus P/O (palmitic acid and oleic acid mixture) treatment induced lipid accumulation in AML12-sh-control cells, while a markedly higher amount of ORO staining was shown in alcohol plus P/O treated AML12-sh-AKR1A1 cells (*p* < 0.001). To determine the effects of AKR1A1 on lipid metabolism, the mRNA levels of fatty acid transport, synthesis, and oxidation were measured. After 48 h alcohol plus P/O treatment, fatty acid transport/synthesis-related genes, including *Cd36, Fasn. Acaca, Cyp4a, Cyp2e1, Lipin1, Ppar-γ*, and *Srebp1* were significantly upregulated (Fig. 6E), and fatty acid oxidation-related genes were significantly downregulated in treated AML12-sh-AKR1A1 cells compared with AML12-sh-control cells (Fig. 6F). To determine the inflammatory effects of AKR1A1 in hepatocytes, LPS-treated AML12 cells were used. After 48 h of treatment, higher *I1-1β* and *Tnf-α*. mRNA levels were detected in treated AML12-sh-AKR1A1 cells than in AML12-sh-control cells (Fig. 6G). To determine the fibrotic effects of AKR1A1 in hepatocyte, the production of extracellular matrix genes in TGF-β1-treated AML12 cells was examined. After 48 h of treatment, ECM mRNA, including *Fn1*, *α-Sma, Colla1*, and *Timp-1*, was significantly upregulated in treated AML12-sh-AKR1A1 cells compared with AML12-sh-control cells (Fig. 6H).

**Figure 6.**
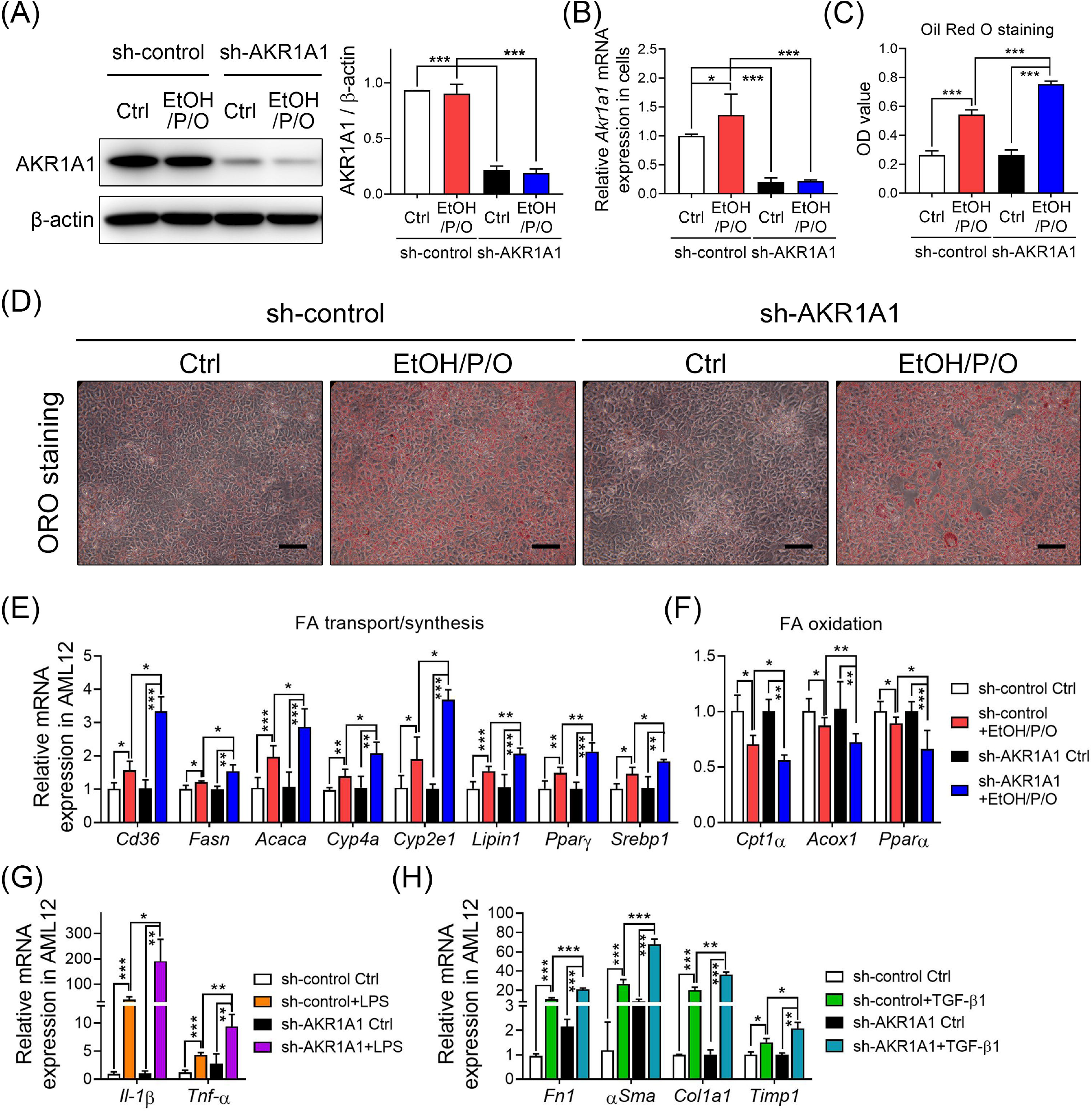
Knockdown of AKR1A1 exacerbates hepatocyte cell line lipid accumulation, inflammation and fibrosis *in vitro*. Knockdown of AKR1A1 exacerbates hepatocyte cell line lipid accumulation, inflammation and fibrosis in vitro. The AKR1A1 knockdown cell line was established by using lentiviral particle transduction. At 48 hours after alcohol plus P/O treatment, (**A**) Western blot analysis and (**B**) qRT–PCR were used to detect AKR1A1 in AML12 cells. (**C**) quantification by extracting intracellular ORO and detecting the absorbance by spectrophotometry at 450 nm and (**D**) Representative images of lipid accumulation were observed by ORO staining. qRT–PCR was performed to detect the mRNA expression levels of (**E**) fatty acid transport/synthesis and (**F**) fatty acid oxidation. (**G**) At 48 h after LPS treatment, qRT–PCR was performed to detect inflammatory genes in AML12 cells. (**H**) At 48 h after TGF-β1 treatment, qRT–PCR was performed to detect fibrotic genes in AML12 cells. Values were normalized to the β-actin gene and expressed in relation to the sh-control group. Data are represented as the mean□±□SD. \**p* < 0.05, \*\**p* < 0.01, and \*\*\**p* < 0.001.

## Discussion

AKR1A1 is known as an enzyme that can detoxify lipid peroxides and cytotoxic products and is also involved in ascorbic acid biosynthesis in different cells under a variety of physiological conditions and pathological challenges. We first found that chronic alcohol feeding increased liver AKR1A1 levels in wild-type mice. The present study provides evidence that hepatic AKR1A1 exerts a protective action against alcohol-induced hepatic oxidative stress, inflammation, steatosis, and fibrosis, possibly through the 4-HNE-p53 signaling pathway.

Chronic alcohol abuse leads to pro-oxidant and antioxidant status imbalance, which may induce programmed cell death in the progression of liver injury [23]. Mounting evidence suggests that the role of AKR1A1 in a variety of cell types is involved in a reduction in the cytotoxicity of aldehyde compounds. The lipid peroxidation aldehyde product 4-HNE, with its high toxicity, has been reported to be involved in the pathogenesis of inflammatory and neurodegenerative disease [24], chronic obstructive pulmonary disease [25], and cancers [26] and in both non-alcoholic fatty liver disease [27] and alcoholic liver disease [28]. As shown in Fig. 3 and Fig. 5, our results demonstrate that deficiency of hepatic AKR1A1 expression in the *Akr1a1*^-/-^ knockout mice could increase 4-HNE levels to exacerbate alcoholic liver associated disease.

Otherwise, AKR1A1 may interact with some anticancer drugs with aldehyde groups, such as doxorubicin, daunorubicin and anthracyclines, to inactivate the drug effects [29,30]. Lacroix et al. [31] found a new role of AKR1A1 with potent prostaglandin F2 (PGF2) synthase activity. The bioactive lipid compound, PGF2 is a member of the PG family with pleiotropic biological functions, which may accelerate the initiation and progression of steatohepatitis, steatosis and advanced fibrosis [32]. A well-known role of AKR1A1 is responsible for ascorbate biosynthesis and action in animals. Taking advantage of genetically deficient mouse models, more latent physiological functions of AKR1A1 have been revealed. Insufficient ascorbate production in AKR1A1-knockout mice leads to further metabolic pathway malfunction, eventually causing orthopedic disorder (osteopenia, spontaneous fractures and osteoporosis) [33,34], impaired formation of spatial memory in juveniles [35], aggressive behaviors [36], embryonic development and neonatal growth disabilities [37], and hepatic steatosis and injury [10–14]. Notably, ascorbate replenishment effectively improves these severe disorders. In this study, we further demonstrate that knockout of *Akr1a1* exacerbates alcohol-induced oxidative stress (Nox-2, MDA, GSSG/GSH ratio, 4-HNE) and apoptosis (caspase 3 and caspase 8) in the livers of mice. Additionally, alcohol can induce higher CYP2E1 levels in the liver, which leads to higher generation of reactive oxygen species (ROS) by metabolizing alcohol. As shown in Fig. 3, high CYP2E1 levels were found in the liver tissue of the alcohol-fed *Akr1a1*^-/-^ mice. In *in vitro* study (Fig. 6), we also demonstrate that knockdown of *Akr1a1* aggravated the effects of alcohol plus P/O-induced oxidative stress and steatosis, LPS-stimulated inflammation, and TGF-β1-induced fibrosis in hepatocyte cell lines.

Tumor protein 53 (p53) is primarily known as a tumor suppressor in response to genotoxic stress in various cancers and has recently been described as a stress sensor that controls a variety of physiological and pathological processes in non-tumor conditions [38]. For example, the toxic lipid peroxidation metabolite 4-HNE is one of the diverse stresses that activates p53, which could induce cell apoptosis [39]. p53 in the liver plays a role in cellular stress response and metabolism to regulate liver homeostasis and dysfunction [40]. High expression levels of hepatic p53 may induce ROS production, cell apoptosis or necrosis, leading to tissue inflammation, which can contribute to subsequent liver steatosis and cirrhosis [41,42]. Taking advantage of genetically deficient mouse models, accumulating evidence has shown that ablation of p53 reduces liver cell apoptosis, oxidative stress, and lipid metabolism in chronic alcohol-[43,44] or non-alcoholic-diet-induced steatohepatitis mouse models [45,46]. As shown in Fig. 5, our results confirm that deficiency of liver AKR1A1 increased phosphorylated p53 levels, which may be involved in the exacerbation of alcoholic liver-associated disease. These data suggest that *Akr1a1* knockout exacerbates ALD, possibly through inhibition of the 4-HNE/p53 signaling pathway in the livers of mice.

Current therapeutic strategies for the treatment of ALD include abstinence-supportive therapies for abstinence, nutrition support, or liver transplantation [47]. In recent years, high-throughput platforms have been widely performed to identify potentially regulated genes or miRNA involved in the progression of alcohol-associated liver disease [48,49]. Restoration of liver function by regulating potentially targeted signaling pathways with compounds or gene therapy approaches has been widely studied [50,51]. Liu et al. [52] replenished fibroblast growth factor 21 in mice to reduce the exacerbation of chronic alcohol-induced hepatic injury. Satishchandran et al. [53] restored liver miR-122 levels by using a gene therapy approach to ameliorate alcoholic liver injury in mice. Stem cell-based therapy in alcohol-associated liver disease showed promising effects through their liver repopulation capacity to drive cell-for-cell replacement, which may restore normal liver function to reduce liver damage and exert immunomodulatory, and anti-fibrotic properties [54–56]. In addition, Brulport et al. [57] mentioned that transplantation of human extrahepatic stem cells to mouse liver showed partial transdifferentiation and horizontal gene transfer.

## Conclusion

In this study, we report a novel finding that knockout of AKR1A1 leads to exacerbation of liver oxidative stress, inflammation, steatosis, and fibrosis in chronic alcohol-fed mice, possibly through elevated 4-HNE production and activated p53 signaling. Therefore, elevating AKR1A1 function may act as a potential therapeutic target for prevention or treatment of alcoholic-associated liver disease.

## Abbreviations

4-HNE: 4-hydroxynonenal
ADH: alcohol dehydrogenase
ADRP: adipose differentiation-related protein
AF: alcohol-fed
AKR1A1: aldo-keto reductase family 1 member A1
ALD: alcoholic-associated liver disease
ALT: alanine aminotransferase
AST: aspartate aminotransferase
Casp3: caspase 3
Casp8: caspase 8
Col1a1: collagen type I alpha 1
Ctgf: connective tissue growth factor
CYP2E1: cytochrome p450 family 2 subfamily E member 1
FASN: fatty acid synthase
GSH: reduced glutathione
GSSG: oxidized glutathione
H&E: hematoxylin and eosin
IHC: immunohistochemistry
IL-1β: interleukin-1β
MDA: malondialdehyde
Nox-2: NADPH oxidase 2
Nqo-1: NAD(P)H quinone dehydrogenase 1
ORO: oil red O
pACC: phospho-acetyl CoA carboxylase
PF: pair-fed
qRT–PCR: quantitative real-time reverse transcription polymerase chain reaction
Sod-1: superoxide dismutase type 1
SREBP1: sterol regulatory element-binding transcription factor 1
Tgf-β1: transforming growth factor beta 1
TNF-α: tumor necrosis factor-α
WT: wild type.

## Authors’ contributions

Conceptualization, Y.-W.L., W.-R.C., K.-Y.C. and C.-M.C.; methodology, C.-C.Y., K.-Y.C. and H.C.F.; validation, Y.-W.L., Y.-C.C. and C.-C.Y.; formal analysis, Y.-W.L., W.-R.C. and H.C.F.; investigation, Y.-W.L. and W.-R.C.; resources, C.-M.C.; data curation, C.-C.Y. and M.-S.C.; writing—original draft preparation, Y.-W.L.; writing—review and editing, C.-M.C.; supervision, C.-M.C.; project administration, Y.-W.L.; funding acquisition, C.-M.C. All authors have read and agreed to the published version of the manuscript.

## Data availability statement

All data generated or analyzed during the current study are available from the corresponding author on reasonable request.

## Acknowledgments

We thank our colleague, Dr. Gary Ro-Lin Chang, in the Molecular Embryology & DNA Methylation Laboratory for his help with discussions and technical issues.

## Supplementary Information

Supplementary data to this article can be found online.

## Additional files

Supplementary **Figure S1**. Effects of a Lieber-DeCarli liquid diet containing 5% alcohol (AF) or maltodextrin (PF) for 8 weeks on C57BL/6 and ICR mice. Supplementary **Table S1**. Primer sequences for qRT-PCR. Supplementary **Table S2.** Primary antibodies used for western blot (WB) and immunohistochemistry analyses. Supplementary **Table S3**. Change in body weight, liver weight and epididymal fat weight in ICR wild-type (WT) or *Akr1a1*^-/-^ mice treatment with pair-fed or alcohol-fed for 8 weeks.

## Declarations

### Ethics approval

All animal experiments were approved by the Institutional Animal Care and Use Committee of National Chung Hsing University (IACUC No. 109-148). Experimental protocols and animal care were provided according to the guideline by National Chung Hsing University, and conformed to the Declaration of Helsinki.

### Consent to participate

Not applicable.

### Consent for publication

Not applicable.

