## Supplementary Table and Figure for "Aldo-Keto Reductase Family 1 Member A1 (AKR1A1) Deficiency Exacerbates Alcohol-Induced Hepatic Oxidative Stress, Inflammation, Steatosis, and Fibrosis"

### Supplementary Figure S1.

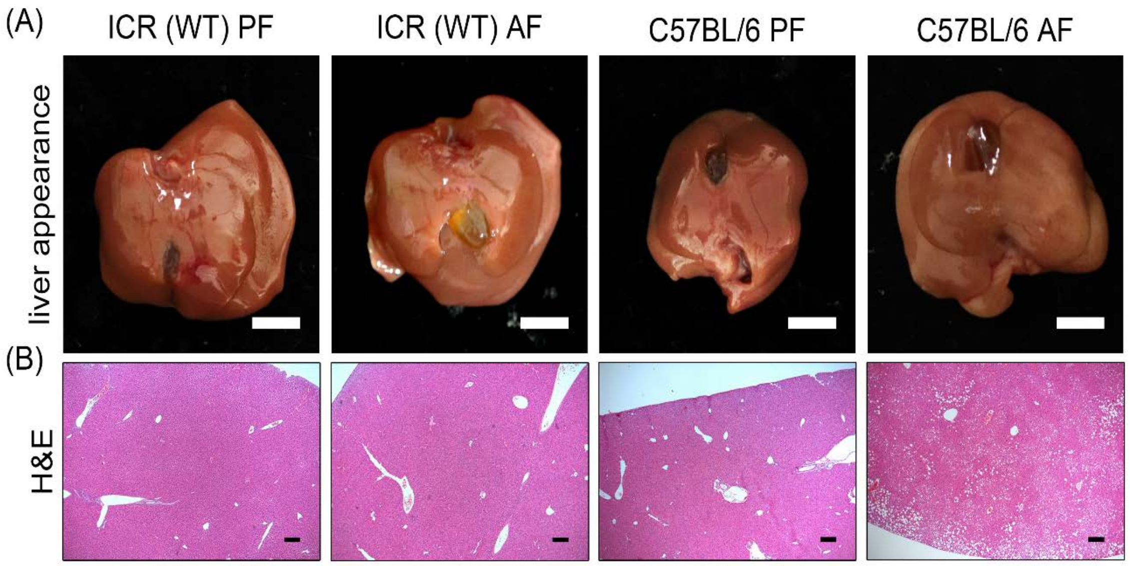


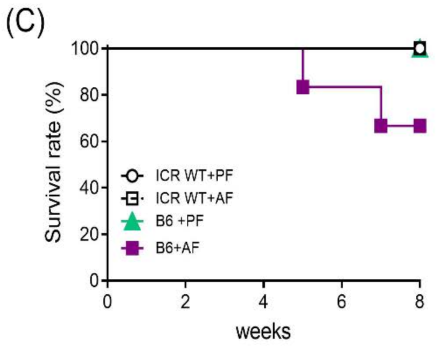


(D)

| | Body weight (g) | Liver weight (g) | Liver weight to body weight ratio (%) |
|---|---|---|---|
| ICR (WT) PF | 34.98 ± 2.49 | 1.55 ± 0.10 | 4.44 ± 0.42 |
| ICR (W)T AF | 33.73 ± 1.66 | 1.57 ± 0.06 | 4.67 ± 0.33 |
| C57BL/6 PF | 27.92 ± 1.08 | 1.04 ± 0.12 | 3.74 ± 0.51 |
| C57BL/6 AF | 24.75 ± 1.05*** | 1.25 ± 0.10** | 5.05 ± 0.44*** |

**Supplementary Figure S1. Effects of a Lieber-DeCarli liquid diet containing 5% alcohol (AF) or maltodextrin (PF) for 8 weeks on C57BL/6 and ICR mice.** (**A**) Gross liver appearance, scale bar = 1 cm, and (**B**) H&E stained liver sections of PF or AF mice for 8 weeks. Scale bar = 200 μm. (**C**) Comparison of survival rates among the four groups. (**D**) Body weight, liver weight, and the percentage of liver to body weight for each group. PF, pair-fed; AF, alcohol-fed; H&E, hematoxylin and eosin.

**Supplementary Table S1. Primer sequences for qRT-PCR**

| Target | Sense | Antisense |
|---|---|---|
| IL-1β | 5'-GCTCATCTGGGATCCTCTCC-3' | 5'-CCTGCCTGAAGCTCTTGTTG-3' |
| TNF-α | 5'-CCCTCACACTCAGATCATCTTCT-3' | 5'-GCTACGACGTGGGCTACAG-3' |
| NOX-2 | 5'-AGTGCGTGTTGCTCGACAA-3' | 5'-GCGGTGTGCAGTGCTATCAT-3' |
| SOD-1 | 5'-TATGGGGACAATACACAAGGCT-3' | 5'-CGGGCCACCATGTTTCTTAGA-3' |
| NQO-1 | 5'-AGAGAGTGCTCGTAGCAGGAT-3' | 5'-GTGGTGATAGAAAGCAAGGTCTT-3' |
| Casp3 | 5'-CTGACTGGAAAGCCGAAACTC-3' | 5'-CGACCCGTCCTTTGAATTTCT-3' |
| Casp8 | 5'-CAACTTCCTAGACTGCAACCG-3' | 5'-TCCAACTCGCTCACTTCTTCT-3' |
| CD36 | 5'-ATGGGCTGTGATCGGAACTG-3' | 5'-TTTGCCACGTCATCTGGGTTT-3' |
| VLDLR | 5'-TGATTGCGAAGACGGTTCTGA-3' | 5'-CCAGGACACGGGGATACACT-3' |
| FATP1 | 5'-CGCTTTCTGCGTATCGTCTG-3' | 5'-GATGCACGGGATCGTGTCT-3' |
| LPL | 5'-GGGAGTTTGGCTCCAGAGTTT-3' | 5'-TGTGTCTTCAGGGGTCCTTAG-3' |
| CPT1α | 5'-AGATCAATCGGACCCTAGACAC-3' | 5'-CAGCGAGTAGCGCATAGTCA-3' |
| ACOX1 | 5'-TAACTTCCTCACTCGAAGCCA-3' | 5'-AGTTCCATGACCCATCTCTGTC-3' |
| PPAR-α | 5'-GCAGCTCGTACAGGTCATCA-3' | 5'-CTCTTCATCCCCAAGCGTAG-3' |
| ACACA | 5'-AATGAACGTGCAATCCGATTTG-3' | 5'-ACTCCACATTTGCGTAATTGTTG-3' |
| FASN | 5'-GGCTCTATGGATTACCCAAGC-3' | 5'-CCAGTGTTCGTTCCTCGGA-3' |
| SREBP1 | 5'-GATCGCAGTCTGAGGAGGAG-3' | 5'-GATAGCAGGATGCCAACAGC-3' |
| LIPIN1 | 5'-CATGCTTCGGAAAGTCCTTCA-3' | 5'-GGTTATTCTTTGGCGTCAACCT-3' |
| CYP2E1 | 5'-GGACCTTTCCCAATTCCTTTCTT-3' | 5'-TCTTGTGGTTCAGTAGCACCT-3' |
| CYP4A | 5'-TTCCCTGATGGACGCTCTTTA-3' | 5'-GCAAACCTGGAAGGGTCAAAC-3' |
| PPAR-γ | 5'-TGTGGGGATAAAGCATCAGGC-3' | 5'-CCGGCAGTTAAGATCACACCTAT-3' |
| TGF-β1 | 5'-CTTCAATACGTCAGACATTCGGG-3' | 5'-GTAACGCCAGGAATTGTTGCTA-3' |
| COL1A1 | 5'-CTGACTGGAAGAGCGGAGAGTAC-3' | 5'-ACAGACGGCTGAGTAGGGAACA-3' |
| CTGF | 5'-ACCTGGAGGAAAACATTAAGAAGG-3' | 5'-AGCCCTGTATGTCTTCACACTG-3' |
| α-SMA | 5'-TCCTTCGTGACTACTGCCGAGC-3' | 5'-AATGGTGATCACCTGCCCGTC-3' |
| FN1 | 5'-TTCAAGTGTGATCCCCATGAAG-3' | 5'-CAGGTCTACGGCAGTTGTCA-3' |
| TIMP1 | 5'-GCAACTCGGACCTGGTCATAA-3' | 5'-CGGCCCGTGATGAGAAACT-3' |
| AKR1A1 | 5'-CAACTGGAGTATTTGGACCTC-3' | 5'-GACATCATCAATCTGCCGAC-3' |
| β-actin | 5'-CGCCACCAGTTCGCCATGGA-3' | 5'-TACAGCCCGGGGAGCATCGT-3' |

Abbreviations: IL-1β, interleukin-1β; TNF-α, tumor necrosis factor-α; Nox-2, NADPH oxidase 2; Sod-1, superoxide dismutase type 1; Nqo-1, NAD(P)H quinone dehydrogenase 1; Casp3, caspase 3; Casp8, caspase 8; VLDLR, very low density lipoprotein receptor; FATP1, fatty acid transport protein 1; LPL, lipoprotein lipase; CPT1α, carnitine palmitoyltransferase 1 alpha; ACOX1, acyl-coA oxidase 1; PPAR-α, peroxisome proliferator-activated receptor alpha; ACACA, acetyl coA carboxylase; FASN, fatty acid synthase; SREBP1, sterol regulatory element-binding transcription factor 1; CYP2E1, cytochrome p450 family 2 subfamily E member 1; CYP4A, cytochrome p450 family 4 subfamily A member; PPAR-γ, peroxisome proliferator-activated receptor gamma; TGF-β1, transforming growth factor beta 1; COL1A1, collagen type I alpha 1; CTGF, connective tissue growth factor; α-SMA, alpha smooth muscle actin; FN1, fibronectin 1; TIMP1, tissue inhibitor of metalloproteinase 1; AKR1A1, aldo-keto reductase family 1 member A1.

**Supplementary Table S2. Primary antibodies used for western blot (WB) and immunohistochemistry analyses**

| Antibody | Dilution used for | | Host | Type | Supplier |
|---|---|---|---|---|---|
| | WB | IHC | | | |
| anti-AKR1A1 | 1:2000 | 1:200 | Rabbit | Polyclonal | Atlas Antibodies AB |
| anti-IL-1β | 1:1000 | - | Mouse | Monoclonal | Cell Signaling Technology |
| anti-TNF-α | 1:2000 | - | Mouse | Monoclonal | Proteintech |
| anti-ADH | 1:5000 | - | Mouse | Monoclonal | Santa Cruz |
| anti-CYP2E1 | 1:2000 | - | Rabbit | Polyclonal | CUSABIO Technology |
| anti-4-HNE | 1:2000 | 1:200 | Rabbit | Polyclonal | Abcam |
| anti-ADRP | 1:500 | - | Rabbit | Polyclonal | Novus Biologicals |
| anti-SREBP1 | 1:500 | - | Rabbit | Polyclonal | Santa Cruz |
| anti-FASN | 1:500 | - | Rabbit | Polyclonal | Santa Cruz |
| anti-p-ACC (Ser79) | 1:1000 | - | Rabbit | Monoclonal | Cell Signaling Technology |
| anti-ACC | 1:2000 | - | Rabbit | Polyclonal | GeneTex |
| anti-p-p53 (Ser15) | 1:1000 | - | Rabbit | Polyclonal | Cell Signaling Technology |
| anti-p53 | 1:1000 | - | Mouse | Monoclonal | Cell Signaling Technology |
| anti-β-actin | 1:10000 | - | Mouse | Monoclonal | Novus Biologicals |

Abbreviations: AKR1A1, aldo-keto reductase family 1 member A1; IL-1β, interleukin-1β; TNF-α, tumor necrosis factor-α; ADH, alcohol dehydrogenase; CYP2E1, cytochrome p450 family 2 subfamily E member 1; 4-HNE, 4-hydroxynonenal; ADRP, adipose differentiation-related protein; SREBP1, sterol regulatory element-binding transcription factor 1; FASN, fatty acid synthase; pACC, phospho-acetyl coA carboxylase.

**Supplementary Table S3. Change in body weight, liver weight and epididymal fat weight in ICR wild-type (WT) or *Akr1a1*$^{-/-}$ mice treatment with pair-fed or alcohol-fed for 8 weeks**

| Parameter | WT+PF | WT+AF | *Akr1a1*$^{-/-}$+PF | *Akr1a1*$^{-/-}$+AF |
|---|---|---|---|---|
| Initial body weight (g) | 32.12 ± 1.39$^{a}$ | 32.35 ± 1.79$^{a}$ | 30.13 ± 3.05$^{a}$ | 26.65 ± 2.95$^{b}$ |
| Final body weight (g) | 35.78 ± 1.82$^{a}$ | 34.54 ± 1.18$^{a}$ | 33.79 ± 2.91$^{a}$ | 25.22 ± 2.60$^{b}$ |
| Liver weight (g) | 1.55 ± 0.10$^{a}$ | 1.57 ± 0.06$^{a}$ | 1.42 ± 0.16$^{b}$ | 1.81 ± 0.10$^{c}$ |
| Liver weight to body weight ratio (%) | 4.34 ± 0.29$^{a}$ | 4.48 ± 0.15$^{a}$ | 4.36 ± 0.20$^{a}$ | 6.41 ± 0.80$^{b}$ |
| Epididymal fat weight (g) | 0.52 ± 0.20$^{a}$ | 0.27 ± 0.09$^{b}$ | 0.40 ± 0.12$^{a}$ | 0.20 ± 0.17$^{c}$ |
| Epididymal fat weight to body weight ratio (%) | 1.33 ± 0.43$^{a}$ | 0.84 ± 0.23$^{b,c}$ | 1.17 ± 0.32$^{a,b}$ | 0.59 ± 0.52$^{d}$ |

The statistical analysis was performed according to Duncan's multiple-range method. Means ± SD followed by the same letters indicate not significant differences between groups but without the same letters indicate significant differences (n=5; $P < 0.05$).
